# MCMBP maintains genome integrity by protecting the MCM subunits from degradation

**DOI:** 10.1101/827386

**Authors:** Venny Santosa, Masato T. Kanemaki

## Abstract

The hetero-hexameric MCM2–7 helicase plays a central role in eukaryotic DNA replication. The expression of MCM2–7 is maintained at a high level for creating dormant origins, which are important for maintaining genome integrity. However, other than transcriptional activation for the de novo synthesis, little is known about how cells maintain a high level of MCM2–7. We show that human MCMBP is a short-lived nuclear protein associating mainly with MCM5, 6, and 7. Loss of MCMBP down-regulates MCM2–7, leading to replication stress and DNA-damage accumulation. Our work demonstrates MCMBP protects the MCM subunits from degradation and suggests its chaperone-like role to achieve a high level of functional MCM2–7 using the nascent and recycled subunits.

## Introduction

The mini-chromosome maintenance 2–7 (MCM2–7) complex is a replicative helicase that plays a central role in eukaryotic DNA replication (Bochman and Schwacha 2009; Masai et al. 2010). Its six subunits are evolutionally related with each other and form a hetero-hexameric complex (Figure S1). In the late M to G1 phase, the MCM2–7 hexamer is loaded onto replication origins bound with ORC1–6, to form pre-replicative complexes (pre-RCs). This reaction, which is also known as licensing, is aided by CDC6 and CDT1. Importantly, the MCM2–7 complex in pre-RCs is inactive as a helicase. Subsequently, upon the activation of two essential kinases, S-CDK and CDC7, MCM2–7 in pre-RCs is converted to the active CDC45-GINS-MCM (CMG) helicase, which becomes the core of the replisome and drives the replication forks in the S phase. The replication forks sometime stall or collapse when they encounter obstacles of both intra- and extra-cellular origins, leading to the accumulation of ‘replication stress’ (Hills and Diffley 2014; Zeman and Cimprich 2014). Failure or incompletion in DNA replication causes genome instability, potentially resulting in cell death, cancer development or genetic disorder. Moreover, cancer cells experience replication stress intrinsically, which is thought to drive cancer evolution (Hills and Diffley 2014). After DNA replication, unused MCM2–7 at dormant origins are removed (Chong et al. 1995; Kubota et al. 1995; Todorov et al. 1995). The meeting of two converging forks during replication termination leads to the ubiquitylation of MCM7 in the CMG helicase, which is recognized and extracted by the p97/CDC48 segregase (Maric et al. 2014; Moreno et al. 2014). The MCM subunits released from chromatin are presumably recycled to form a functional MCM2–7 hexamer for the next round of replication.

Proliferating cells express high levels of MCM2–7, which are loaded onto origins in 3– 10-fold excess over the level used for normal origin firing (Edwards et al. 2002). Only a subset of loaded MCM2–7 at origins are used for DNA replication, with the remaining complexes creating dormant origins, which are used as a back-up when cells are challenged by replication stress. In fact, metazoan cells with reduced licensed origins are sensitive to replication stress and DNA damage (Woodward et al. 2006; Ge et al. 2007; Ibarra et al. 2008). Mice expressing a reduced level of MCM2–7 are prone to developing cancers (Pruitt et al. 2007; Shima et al. 2007; Kunnev et al. 2010; Kawabata et al. 2011). Clearly, the maintenance of a high level of MCM2–7 is important for genome maintenance in proliferating cells. The *MCM* genes are transcriptionally up-regulated by E2F following growth stimulation (Leone et al. 1998; Ohtani et al. 1999).

However, other than transcriptional activation for the de novo synthesis of the MCM subunits, little is known about how a high level of functional MCM2–7 hexamers is maintained in proliferating cells.

The MCM-binding protein (MCMBP) was identified as a protein that associates with the MCM subunits, with the exception of MCM2, and was proposed to form an alternative MCM–MCMBP complex by replacement of MCM2 (Sakwe et al. 2007). However, overexpressed MCMBP can interact with all MCM subunits with different affinity (Nguyen et al. 2012). MCMBP is highly conserved in eukaryotes, except for its absence in budding yeast, and has an MCM-like domain with no ATPase motifs, suggesting that it shares a common ancestor with the MCM proteins (Figure S1). The reported phenotypes stemming from the loss of MCMBP vary among species. Sister-chromatid cohesion was defective in an ETG1/MCMBP mutant of *Arabidopsis thaliana* (Takahashi et al. 2010). Defects in the cell cycle and licensing were found in fission yeast after the inactivation of Mcb1/MCMBP (Ding and Forsburg 2011; Li et al. 2011; Santosa et al. 2013). MCMBP depletion in xenopus egg extracts yielded a defect in the unloading of MCM2–7 from chromatin in the late S phase (Nishiyama et al. 2011). Depletion of MCMBP in *Trypanosoma brucei* caused defects in gene silencing and DNA replication (Kim et al. 2013; Kim 2019). MCMBP depletion in human cells caused nuclear deformation (Jagannathan et al. 2012). However, the primary role of MCMBP remains elusive and it is not known whether the defects observed in these organisms were derived from a single primary role of MCMBP. To understand the primary function of MCMBP in human cells, we adopted a modern genetic approach combined with biochemistry and cytology.

## Results and Discussion

Initially, we fused a 3-tandem FLAG tag to the C terminus of the endogenous MCMBP protein using CRISPR–Cas9 in human colorectal cancer HCT116 cells. To understand how MCMBP associates with the MCM proteins, we prepared an extract from this cell line and fractionated it using size-exclusion chromatography (Figure 1A). MCM2, 4, 6 and 7 peaked at the fractions corresponding to the size of the MCM2–7 hexamer, although these proteins were also detected in smaller fractions. Conversely, MCM3 and 5 peaked between 440 and 150 kDa, suggesting that these two proteins form a smaller subcomplex as previously reported (Kimura et al. 1996). Importantly, MCMBP-FLAG peaked at smaller fractions, similar to that observed for MCM3 and 5. To investigate the association between MCM proteins and MCMBP, we immuno-purified MCMBP-FLAG from the extract and the eluted sample was fractionated according to the same protocol (Figure 1B). We found that a small amount of MCM4 associated with MCMBP-FLAG. Importantly, MCMBP-FLAG associated mainly with MCM5, 6 and 7 in smaller fractions, suggesting that MCMBP associates mainly with MCM5, 6 and 7 monomers and/or subcomplexes containing these subunits.

**Figure 1:**
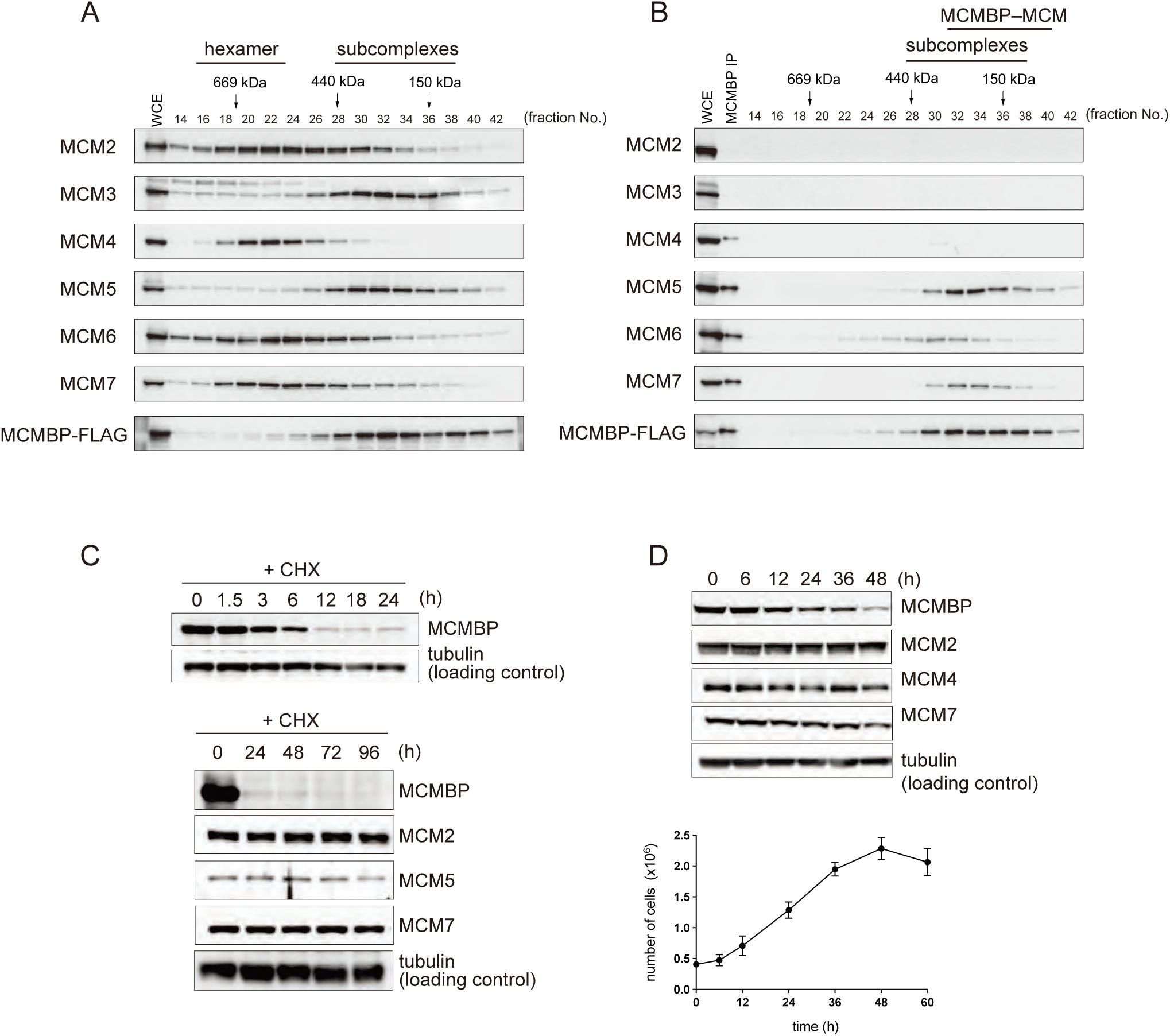
Association between MCMBP and the MCM subunits. (A) WCE prepared from HCT116 cells expressing MCMBP-FLAG was size fractionated for Western blotting. (B) Proteins purified by FLAG IP were size fractionated for Western blotting in a similar manner. Scale bars, 16 μm. (C) HCT116 MCMBP-mAC cells were treated with 25 μM CHX and harvested at the indicated time points. (D) Kinetics of MCMBP expression. HCT116 MCMBP-mAC cells were grown until confluency. The upper panel shows Western blotting data of the indicated proteins. The growth of the cells is shown in the lower panel, indicating that the cells reached confluency at 48 h. Error bars, standard error (n = 3).

The addition of a translation inhibitor, cycloheximide (CHX), revealed that MCMBP was a short-lived protein (t_1/2_ < 3 h) (Figure 1C). Conversely, the half-life of MCM2, 5 and 7 was longer than 72 h in HCT116 cells treated with CHX (note that the proliferation of the treated cells was arrested). Subsequently, we noted that the expression level of MCMBP was controlled by cell proliferation. The expression level of MCMBP decreased after the induction of growth arrest by contact inhibition (Figure 1D). In contrast, the levels of MCM2, 4 and 7 were mostly maintained throughout the time course of the experiment. This notion was also confirmed in cells in which growth was arrested by serum starvation (Figure S1B and C). These results show that MCM2–7 in non-proliferating cells are relatively stable and suggest that MCMBP plays a role that is related to cell proliferation.

To analyse the phenotype of MCMBP-deficient cells, we initially aimed to knockout the *MCMBP* gene in HCT116 cells. However, the possible knockout cells obtained after transfection formed small colonies and did not expand (data not shown). This is consistent with the results obtained for fitness genes, as revealed by a CRISPR-based forward screening (Hart et al. 2015). Therefore, we generated a conditional MCMBP cell line using the auxin-inducible degron (AID) technology (Figure S2A) (Nishimura et al. 2009). We introduced a mini-AID and Clover (mAC) tag at the C-terminus of the endogenous MCMBP in HCT116 parental cells constitutively expressing OsTIR1 from the safe-harbour *AAVS1* locus (Figure S2B) (Natsume et al. 2016). The resulting conditional cells expressed an MCMBP-mAID-Clover fusion protein (MCMBP-mAC) that can be targeted for degradation by the addition of auxin (Figure 2A). MCMBP was mainly localized in the nucleus, as reported previously (Sakwe et al. 2007), and the nuclear signal of MCMBP-mAC had disappeared almost completely 1 day after auxin addition (Figure 2B).

**Figure 2:**
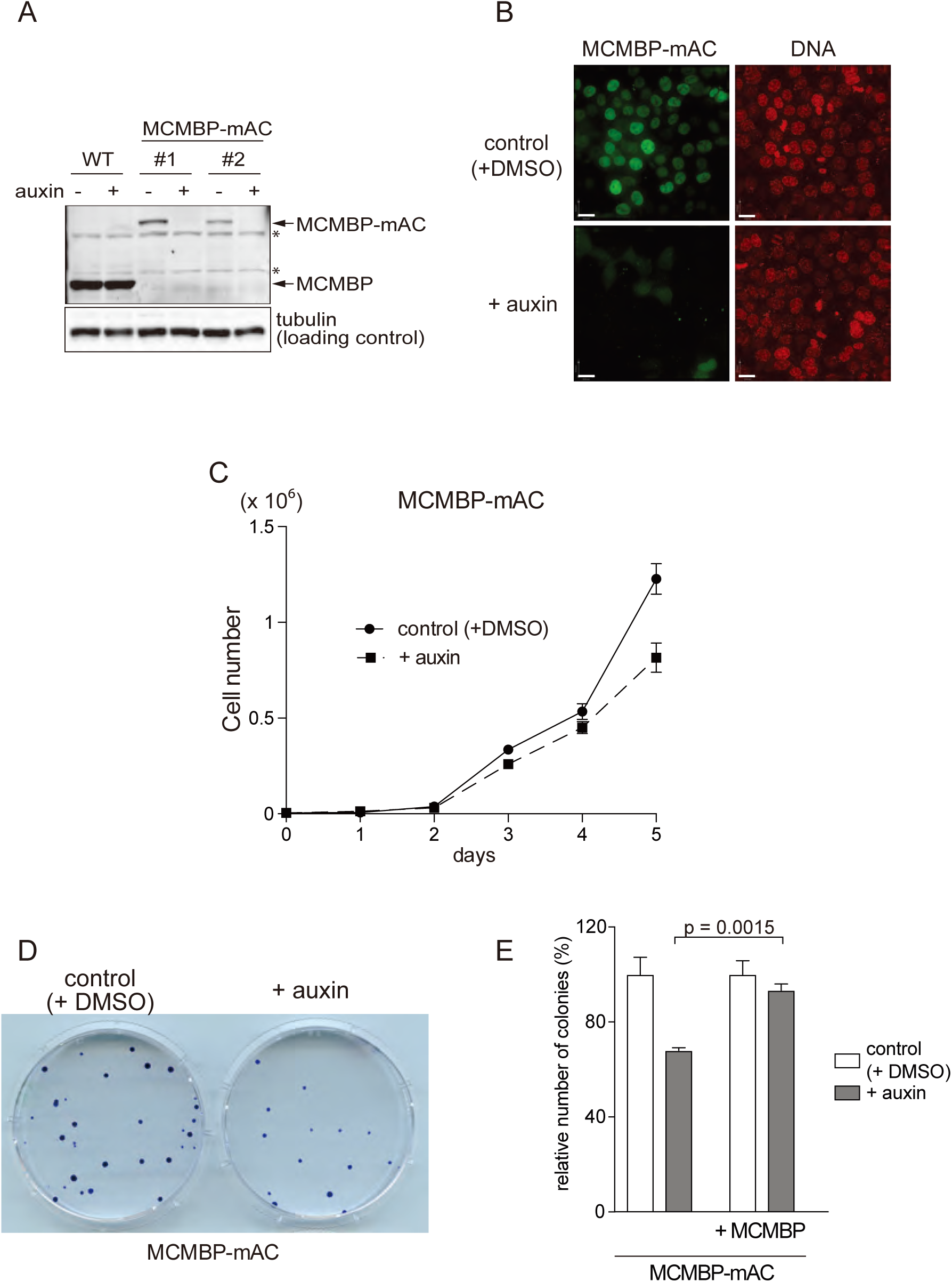
Growth defects after rapid depletion of MCMBP. (A) HCT116 MCMBP-mAC cells were treated with or without 500 μM IAA for 24 h prior to harvest. The asterisks show the position of a non-specific protein. (B) MCMBP depletion detected using microscopy. HCT116 MCMBP-mAC cells were treated with or without 500 μM IAA for 24 h prior to observation. DNA was stained using 0.1 μM SiR-DNA (Spirochrome). Scale bars, 16 μm. (C) HCT116 MCMBP-mAC cells were grown in medium with or without 500 μM IAA. Error bars, standard error (n = 3). (D) One hundred HCT116 MCMBP-mAC cells were seeded in a 10 cm dish and grown in medium with or without 500 μM IAA for 13 days before staining with crystal violet. (E) Colonies were counted. Cells that were rescued by introducing an MCMBP expression construct are presented in the two columns on the right. Error bars, standard error (n = 3).

To test the impact of MCMBP depletion on cell proliferation, we treated the MCMBP-mAC cells with auxin and monitored their growth. The MCMBP-deleted cells showed a growth defect 3 days after auxin addition compared with the mock-treated cells (Figure 2C). We tested colony formation in the presence or absence of auxin and found that MCMBP-depleted cells formed fewer and smaller colonies compared with the mock-treated cells (Figure 2D and E). To confirm that the proliferation defect was caused by MCMBP depletion, we rescued the MCMBP-mAC cells by introducing an MCMBP-expressing construct at the *ROSA26* locus. The number of colonies formed by the rescued cells was significantly increased, even though MCMBP-mAC was depleted by the addition of auxin (Figure 2E). Furthermore, hTERT RPE1 and HeLa cells with depletion of MCMBP by siRNA also showed proliferation defects, similar to MCMBP-depleted HCT116 cells (Figure S2C and D). hTERT PRE1 cells showed a more pronounced defect that might have been caused by p53 activation following DNA-damage accumulation (this will be shown and discussed later). Taken together, these findings led us to conclude that MCMBP is important for the long-term survival of human cells.

Two questions arose at this stage of the research: (1) why did the MCMBP-depleted cells gradually become sick? And (2) what is the relationship between the control of the expression of MCMBP and cell proliferation? MCMBP-mAC was rapidly depleted within 6 h after auxin addition (Figure 3A). Interestingly, we noted that the expression level of MCM2, 4 and 7 was gradually reduced 24 h after auxin addition in MCMBP-depleted cells. We confirmed this notion by microscopical examination of MCM5 and 7 in MCMBP-mAC cells treated with auxin for 5 days (Figure 3B). To clarify whether MCMBP controls the expression of MCM2–7 at the mRNA level, we assessed the mRNA levels of all *MCM* genes (Figure S3). We found that the MCM mRNA levels were not largely affected, supporting the idea that MCMBP maintains MCM expression at the protein level. To generalize our finding that MCMBP maintains the expression of MCM2–7 to other human cell lines, we depleted MCMBP by siRNA in hTERT RPE1 and HeLa cells and found that MCM2, 4, 5 and 7 were also reduced (Figure 3C and D). Based on the fact that down-regulation of one MCM subunit causes degradation of the other MCM subunits (Kawabata et al. 2011) and the observation that MCMBP associates with MCM5, 6 and 7 monomers and/or subcomplexes (Figure 1B), we concluded that MCMBP protects MCM5, 6 and 7 from degradation to maintain the expression of MCM2–7 in proliferating cells.

**Figure 3:**
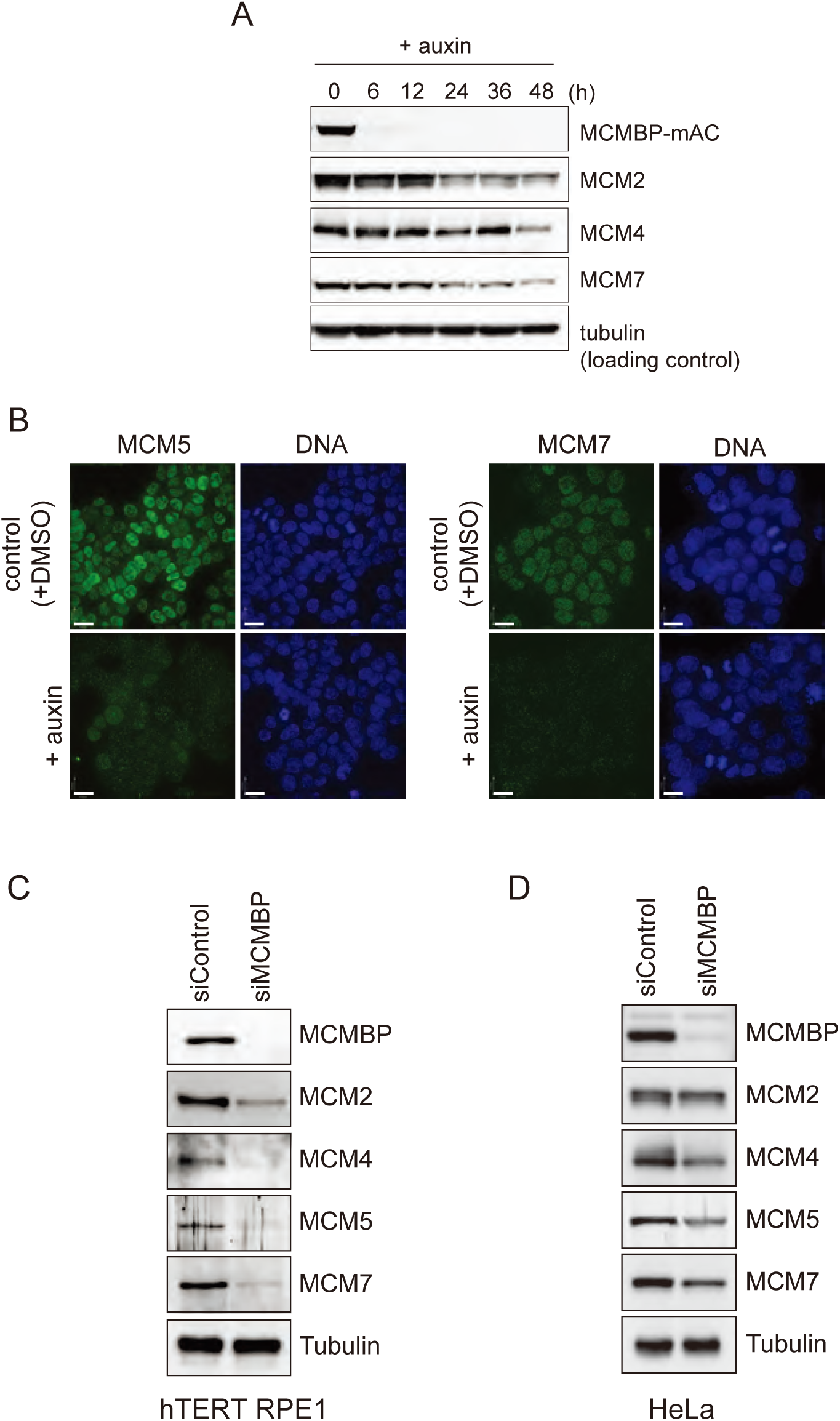
MCMBP depletion down-regulates MCM2–7. (A) HCT116 MCMBP-mAC cells were treated with 500 μM IAA and harvested at the indicated time points. (B) HCT116 MCMBP-mAC cells were treated with or without 500 μM IAA for 5 days before immunostaining, to detect MCM5 and MCM7. DNA was stained with 5 μg/ml of Hoechst33342. (C) hTERT RPE-1 and HeLa cells were treated with siMCMBP or siControl for 3 days prior to harvest. Protein samples were separated and transferred for Western blotting using the indicated antibodies.

MCM2–7 are excessively expressed in proliferating cells, which is important for maintaining genome integrity, in particular when cells experience replication stress (Woodward et al. 2006; Ge et al. 2007; Ibarra et al. 2008). Because MCMBP-depleted cells showed reduced expression of MCMs, we expected that the depleted cells would experience replication defects caused by a reduced number of licensed origins. Initially, we investigated origin licensing by looking at chromatin-bound MCM2–7 in the G1 phase. For this purpose, the MCMBP-mAC cells were treated with or without auxin before G1 arrest induced by lovastatin (Figure 4A and B). G1-arrested cells were lysed and chromatin-bound and -unbound fractions were blotted. Interestingly, MCMBP did not associate with chromatin, unlike MCMs (Figure 4C, – auxin Chr). Consistent with the results shown in Figure 3A, MCMBP depletion reduced the expression level of MCMs in whole-cell extracts (Figure 4C, + auxin WCE), resulting in reduced chromatin association of MCM2, 4 and 7 in G1-arrested cells (Figure 4C, compare the Chr lane of –/+ auxin samples). This result indicates that the number of licensed origins is reduced in MCMBP-depleted cells. Subsequently, we investigated cell-cycle progression in MCMBP-depleted cells by flow cytometry and found that the depleted cells accumulated in the late S to G2 phases (Figure 4D). We recapitulated this phenotype by reducing MCM2 or MCM5 expression by siRNA while maintaining the expression level of MCMBP (Figure S4A and B). These results strongly suggest that MCMBP-depleted cells experience DNA replication defects because of reduced origin licensing. Defects in DNA replication cause under-replication, leading to the generation of DNA lesions in the following G1 phase (Lukas et al. 2011). We confirmed the accumulation of DNA lesions in MCMBP-depleted cells by monitoring 53BP1 nuclear bodies (Figure 4E). A study using xenopus egg extracts showed that MCMBP is involved in MCM2–7 removal from chromatin DNA (Nishiyama et al. 2011). To test whether this was also the case in human cells, the MCMBP-mAC cells were treated with or without auxin before G2 arrest by a CDK inhibitor, RO-3306 (Figure S4C and D). The G2-arrested cells were lysed and the chromatin association of MCMs was investigated. We found that chromatin-bound MCM2, 5 and 7 were down-regulated in MCMBP-depleted cells because of the reduced expression of MCM2–7 (Figure S4E). Because we did not observe accumulation of chromatin-bound MCMs, we concluded that MCMBP is not important for MCM2–7 removal from chromatin DNA in human cells.

**Figure 4:**
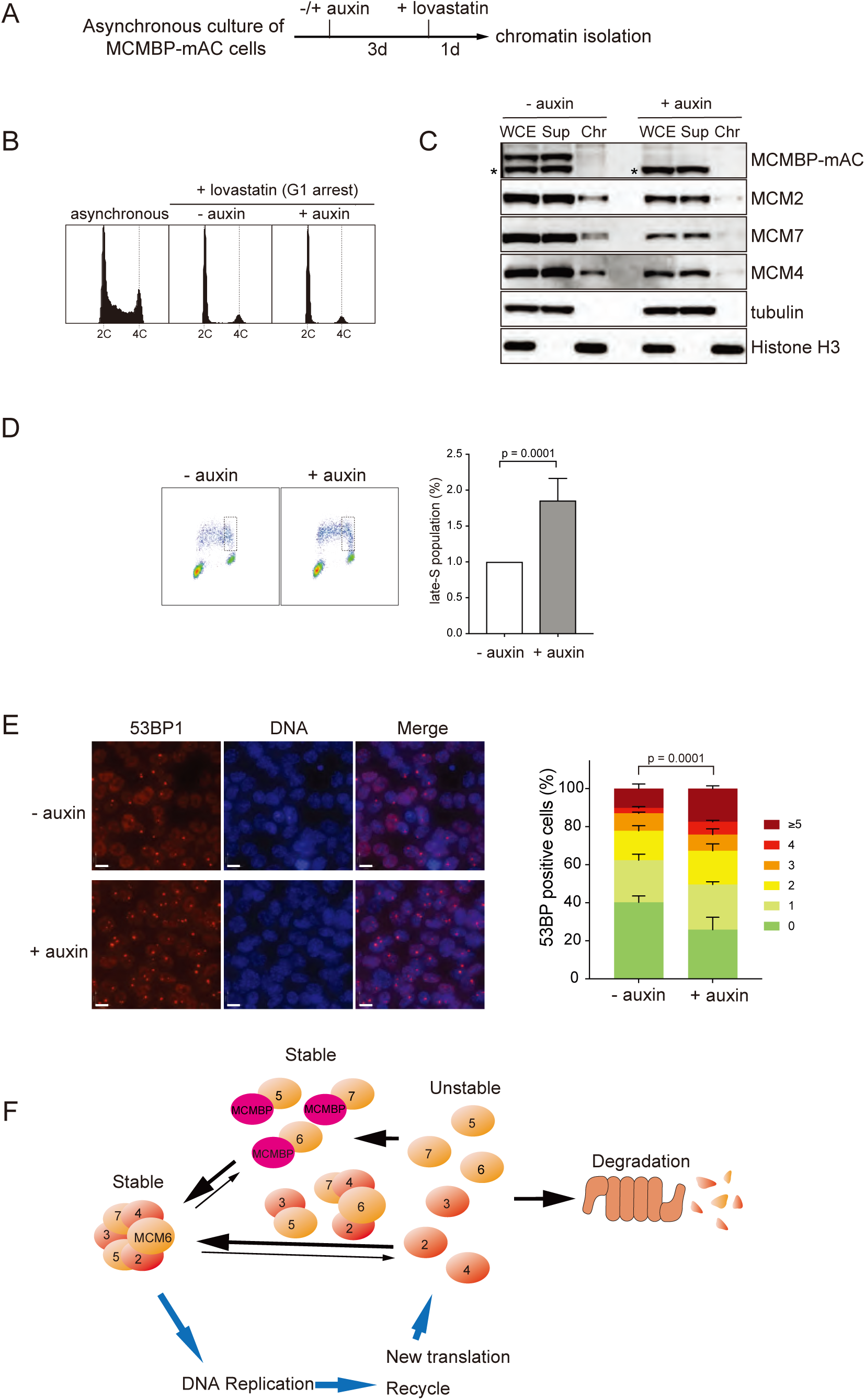
Loss of MCMBP causes a replication defect leading to genome instability. (A) Experimental scheme used to test the chromatin association of MCMs in the G1 phase. (B) Flow cytometry to confirm G1 arrest by lovastatin. (C) G1-arrested cells treated with or without 500 μM IAA were fractionated for Western blotting using the indicated antibodies. Note that MCMBP did not associate with chromatin (– auxin, Chr). (D) HCT116 MCMBP-mAC cells were treated with or without 500 μM IAA for 3 days. In the last 30 min of the incubation, 30 μM BrdU was added (prior to harvest). After staining with an anti-BrdU antibody and PI, the cells were analysed by flow cytometry. The quantification of late-S-phase cells is shown on the right. Error bars, standard error (n = 3). (E) HCT116 MCMBP-mAC cells were treated with or without 500 μM IAA for 5 days. The cells were harvested and stained with an anti-53BP1 antibody. Representative images are shown on the left. Scale bars, 11 μm. The quantification of 53BP1 nuclear bodies is presented on the right. Error bars, standard error (n = 3). (F) A schematic model showing the function of MCMBP in human cells. The MCM2–7 hexamer is stable. After DNA replication, chromatin-bound MCM2–7 complexes are released. Released and newly synthesized MCM subunits are fragile and must be protected by MCMBP or by hexamer formation. MCMBP mainly associates with MCM5, 6 and 7 and protects them from degradation. MCMBP might promote the nuclear import of associating MCM subunits and/or hexamer formation. Loss of MCMBP leads to the degradation of unstable MCM subunits, thus leading to reduced expression of the MCM2–7 hexamer.

Our study demonstrated that MCMBP associated weakly with MCM4 and associated mainly with MCM5, 6 and 7 monomers and/or subcomplexes, thus protecting them from degradation (Figure 4F). This protective function is important to support the high expression of MCM2–7 for maintaining genome integrity in proliferating cells. It appears that the MCM2–7 hexamer is stable in non-proliferating cells (Figures 1D and S1C). Conversely, the MCM subunits are highly synthesized in proliferating cells (Leone et al. 1998; Ohtani et al. 1999). Moreover, unused MCM2–7 on chromatin are removed via an unknown mechanism (Chong et al. 1995; Kubota et al. 1995; Todorov et al. 1995) and the CMG helicase (CDC45–MCM–GINS) is also removed after MCM7 ubiquitylation by the p97 segregase (Maric et al. 2014; Moreno et al. 2014). We propose that MCMBP is a chaperone that protects the de novo synthesized and recycled MCM subunits from degradation (Figure 4F).

Other than protecting the MCM subunits from degradation, what is the molecular function of MCMBP? We reported previously that the nuclear localization of MCM2–7 was lost in fission yeast after the inactivation of Mcb1/MCMBP (Santosa et al. 2013). We did not observe an obvious re-localization of MCM2–7 to the cytoplasm after MCMBP depletion in the human cells tested here (data not shown). However, we do not rule out the possibility that MCMBP helps transport nascent MCM subunits from the cytoplasm to the nucleus. Indeed, MCMBP contained a bipartite NLS (Figure S1A). Intriguingly, only MCM2 and 3 contained clear monopartite NLSs, whereas MCM4, 5, 6 and 7, which interacted with MCMBP, did not have a predicted NLS (Figures 1B, S1A) (Kimura et al. 1996). In the nucleus, MCMBP might help to assemble the MCM2– 7 hexamer. FKBP51, a co-factor for HSP90, associates with MCMBP independently of HSP90 (Taipale et al. 2014). FKBP51 displays peptidyl-prolyl isomerase activity, which helps to promote structural changes in its substrates (Hahle et al. 2019). We hypothesize that MCMBP acts together with FKBP51 to promote the formation of MCM2–7 hexamers in the nucleus. Consistent with this hypothesis, MCMBP did not associate with chromatin (Figures 4C and S4E), suggesting that MCMBP functions mainly in the nucleoplasm. The MCM2–7 hexamer is reportedly disrupted by the excess addition of MCMBP in vitro (Nishiyama et al. 2011; Nguyen et al. 2012; Kusunoki and Ishimi 2014). We also found that overexpression of MCMBP caused growth defects in human cells (data not shown). These results suggest that MCMBP promotes both the assembly and disassembly of the MCM2–7 hexamer in the nucleoplasm as a chaperone, and that the balance between MCMBP and the MCM subunits is important for maintaining the pool of functional MCM2–7 hexamers (Figure 4F). This hypothesis is supported by the observation that the expression level of MCMBP was down-regulated in non-proliferating cells, in which the synthesis of the MCM subunits is attenuated, whereas the expression of MCM2–7 was stably maintained (Figures 1D and S1C) (Leone et al. 1998; Ohtani et al. 1999). We and others noted that MCMBP also associates with MCM8 and 9, which form the MCM8– 9 hexamer (data not shown) (Kusunoki and Ishimi 2014). We predict that MCMBP is also important for maintaining the expression of MCM8–9 for homologous recombination repair in mitosis and meiosis (Lutzmann et al. 2012; Nishimura et al. 2012; Park et al. 2013).

MCMBP is highly conserved (32% identical and 50% positives between humans and *Arabidopsis*); therefore, we expect that MCMBP has a conserved function. We did not observe reduced expression of MCMs in fission yeast with inactivated MCMBP/Mcb1 (Santosa et al. 2013). However, it is possible that the formation of the MCM2–7 hexamer was perturbed, causing re-localization of the MCM monomers or subcomplexes to the cytoplasm in fission yeast. Similarly, the reported defects in nuclear deformation in human cells, in sister-chromatid cohesion in plants, in gene silencing in *Trypanosoma brucei* and in the cell cycle in fission yeast might have been caused by a reduction in the amount of the functional MCM2–7 hexamer after loss of MCMBP (Takahashi et al. 2010; Ding and Forsburg 2011; Li et al. 2011; Jagannathan et al. 2012; Kim et al. 2013).

High expression of MCMBP is associated with an increase in risk of relapse of leukaemia and colon and head-and-neck carcinomas after treatment (Quimbaya et al. 2014). A similar trend was reported for pancreatic and renal cancers (the Human Protein Atlas; https://www.proteinatlas.org/ENSG00000197771-MCMBP/pathology). Based on the fact that excessive MCM2–7 loading confers resistance against replication stress (Woodward et al. 2006; Ge et al. 2007; Ibarra et al. 2008), the up-regulation of MCMBP likely helps cancer survival by increasing the expression of MCM2–7. Therefore, MCMBP may be a potential target that can be combined with cancer chemotherapy in the future.

## Materials & Methods

### Plasmids

To insert a 3-tandem FLAG or mAID–Clover cassette at the C-terminal coding region of the *MCMBP* gene, we designed and generated a CRISPR plasmid targeting the 5′– GTTTGCCCATTACTCTT/CAT(PAM:AGG)–3′ sequence using pX330-U6-Chimeric_BB-CBh-hSpCas9 (Addgene #42260, gift from Dr. Feng Zhang), according to a published protocol (Ran et al. 2013). The donor plasmids were based on pMK283, 286, 286 and 290 (Yesbolatova et al. 2019). To construct donor plasmids, we synthesized 200 bp homology arms up- and down-stream of the targeting site, and a 3-tandem FALG or mAID–Clover cassette with a neomycin or hygromycin selection marker was cloned (Natsume et al. 2016). The rescue construct for expressing MCMBP was based on pMK404, which contains CMV promoter–multi-cloning sites– SV40 poly and a blasticidin selection marker flanked by homology arms for *ROSA26* insertion. The full-length *MCMBP* coding region was cloned at the cloning site. For insertion at the *ROSA26* locus, a CRISPR plasmid targeting the 5′– TTGCAGCTCGCGCCGGT/TTT(PAM:TGG)–3′ was prepared similarly to the MCMBP CRISPR plasmid. All plasmids described in this paper are available from Addgene.

### Cell culture

HCT116, hTERT RPE1 and HeLa cells were cultured according to the standard protocol provided by ATCC. To edit genomic DNA in HCT116 cells, CRISPR and donor plasmids were transfected and clones were selected as described previously (Yesbolatova et al. 2019). Insertion was confirmed by genomic PCR and the expression of tagged proteins was subsequently confirmed by Western blotting. To induce degradation of MCMBP-mAC, the cells were treated with 500 μM IAA. To knockdown proteins by RNAi, cells were seeded on a 6-well plate and transfected with 10 pmol of siControl (negative control #4390843, Ambion Silencer Select), siMCM2 (#s8536, Ambion Silencer Select), siMCM5 (Ge et al. 2007), or siMCMBP (#s36586, Ambion Silencer Select) using Lipofectamine RNAiMAX (ThermoFisher Scientific) according to the manufacturer’s protocol. The transfected cells were cultured for 3 days prior to analysis.

### Protein extraction and Western blotting

Cells were collected using trypsin-EDTA and washed with D-PBS(–). Proteins were extracted by adding RIPA buffer (25 mM Tris-HCl, pH 7.6, 150 mM NaCl, 1% NP-40, 1% sodium deoxycholate, 0.1% SDS and 1× cOmplete-EDTA free). After suspension and clarification by centrifugation, the supernatant was collected. Protein concentration was checked using the Bradford reagent (Bio-Rad, #5000006) on a Smartspec 3000 instrument (Bio-Rad). SDS sample buffer (125 mM Tris-HCl, pH 6.8, 4% SDS, 20% glycerol, 0.01% bromophenol blue and 10% 2-mercaptoethanol) was added and samples were boiled for 5 min at 95°C. Proteins were separated on SDS–PAGE and transferred to a Hybond ECL membrane (GE Healthcare) using Trans-Blot Turbo (Bio-Rad). After blocking with 1% skim milk in TBST for 1 h at room temperature, the membrane was blotted using an appropriate antibody. Detection was performed using a chemiluminescence system with the Amersham ECL Prime reagent (GE Healthcare) or using a fluorescence-based system with ChemiDoc MP (Bio-Rad).

### Antibodies

Listed in Table S1.

### Biochemical experiments

To fractionate cell extracts, cells were harvested and washed with D-PBS(–) before resuspension in cytoskeletal (CSK) buffer (10 mM PIPES-KOH, pH 6.8, 100 mM NaCl, 300 mM sucrose, 1 mM EGTA and 1 mM MgCl_2_) supplemented with 1 mM DTT, 1× cOmplete EDTA-free (Sigma-Aldrich) and 0.5% Triton X-100. The suspension was kept on ice for 20 min and vortexed periodically, to allow extraction. Whole-cell extracts were collected at the end of the reaction and the remaining sample was centrifuged at 5000 rpm for 5 min at 4°C, to separate the supernatant (soluble fraction) from the pellet (chromatin fraction). SDS sample buffer was then added to each fraction and the samples were boiled at 95°C for 5 min. For size fractionation using a Superose6 column, 1.5 × 10^8^ cells were harvested and washed twice with D-PBS(–) before re-suspension in IP buffer (20 mM HEPES-KOH, pH 7.4, 150 mM NaCl, 1 mM EDTA, 0.5% NP-40, 1 mM ATP and 5% glycerol) supplemented with 0.3× PhosStop (Sigma-Aldrich) and 1× cOmplete EDTA-free. The suspension was sonicated (80% duty cycle, 1.0 power, 5 s) using a Branson Sonifier 250 instrument. MgCl_2_ and benzonase (Sigma-Aldrich) were added to the final concentrations of 5 mM and 150 U, respectively. The suspension was kept on ice for 30 min, with occasional inversion. The suspension was subsequently centrifuged at 5,600 rpm for 5 min at 4°C and the supernatant was transferred to a new tube. This clarification process was repeated at a centrifugation speed of 13,000 rpm. The cell extract was then filtrated using a 0.45 μm Spin-X centrifuge tube filter (Costar). At this point, 3/4 of the total cell extract was separated for immunoprecipitation, while the remainder of the extract was fractionated using Superose6 10/300GL (GE Healthcare) fitted to an AKTA explorer system (GE Healthcare) in isocratic-elution manner with a fraction size of 200 μl. The collected fractions were mixed with SDS sample buffer and boiled at 95°C for 5 min. The remaining 3/4 of the total cell extract was subjected to immunoprecipitation. Antibody-conjugated bead preparation was performed as follows: 120 μl of DynaBeads Protein G (ThermoFisher Scientific) was transferred to a tube and washed twice with IP buffer. For conjugation, 800 μl of PBST (PBS with 0.05% Tween-20) and 12 μl of the FLAG M2 antibody (Sigma-Aldrich) were added to the beads and rotated at room temperature for 30 min. The coupled beads were washed with IP buffer once and blocked using 500 μl of 1% BSA in PBS for 30 min with rotation at room temperature. The beads were then rinsed twice with IP buffer before incubation with the cell extract. The mixture was rotated at 4°C for 2 h and the beads were rinsed with IP buffer containing 300 mM NaCl three times, with 5 min of rotation each time. Proteins were eluted using 500 μl of IP Buffer containing 0.25 mg/ml of the 3×Flag peptide (Sigma-Aldrich) and 1× cOmplete EDTA-free for 15 min at 4°C. The elution was fractionated using Superose6 10/300GL.

### Cell-cycle synchronization

For synchronization at the G1 phase, asynchronously growing cells were treated with 20 μM lovastatin (LKT Laboratories) for 24 h. To induce G2 arrest, asynchronously growing cells were treated with 9 μM RO-3306 for 16 h.

### Flow cytometry

To analyse DNA content, cells were fixed overnight using 70% ethanol. After the removal of the ethanol and subsequent wash with PBS, the cells were re-suspended and stained with a propidium iodide (PI) solution (50 μg/ml of RNaseA, 40 μg/ml of PI and 1% BSA) for 30 min at 37°C. Cells treated with BrdU for 30 min prior to harvesting were fixed using 95% ethanol overnight. BrdU staining was performed as follows: after the removal of ethanol and subsequent washing with PBS, the cells were treated with 2 M HCl containing 0.5% Triton X-100 for 30 min. To quench denaturing reactions and calibrate pH, the cells were incubated in 0.1 M Na_2_B_4_O_7_ solution for 30 min to 1 h. Subsequently, the cells were washed using Antibody Solution (1% BSA/PBS containing 0.2% Tween-20). A mouse monoclonal anti-BrdU antibody (BD Biosciences) was added to the antibody solution at 100× dilution. For labelling, the reaction was allowed to proceed for 1 h with gentle rotation. After washing, Antibody Solution containing 1/100× FITC-conjugated goat polyclonal anti-mouse IgG (Jackson Laboratory) was added to the cells, which were incubated for 1 h with gentle rotation. To remove non-specific binding, the cells were washed twice with Antibody Solution. DNA was subsequently stained with PI, as described above. All samples were filtered using a 42 μm nylon mesh cell strainer (Falcon) prior to analysis on a BD Accuri C6 flow cytometer (BD Biosciences). The acquired data were analysed using FCS Express 4 Flow Cytometry Edition (De Novo Software) and the FlowJo software.

### Microscopy

For immunostaining, cells were cultured in 35 mm glass-bottom culture dishes (MatTek) in supplemented McCoy’s 5A medium without Phenol Red (Gibco). Cells were fixated with 3.7% formaldehyde in PBS for 15 min, permeabilized with 0.5% TritonX-100 in PBS for 20 min, and blocked with 3% skim milk in PBS-T (0.05% TritonX-100 in PBS) for 1 h. After blocking, the appropriate primary antibody (diluted 500× in 1% BSA/PBS) was added onto the coverslip and incubated for 1 h, followed by incubation with the appropriate secondary antibody (diluted 1000× in 1% BSA/PBS), in a similar manner. DNA was stained with 5 μg/ml of Hoechst33342 (Life Technologies) for 30 min. All steps were performed at room temperature. Vectashield Mounting Medium (Vector Labs) was then added onto the coverslip. All microscopy was performed using a Delta Vision deconvolution microscope (GE Healthcare).

### RNA detection

HCT116 MCMBP-mAC cells were seeded on 10 cm dishes and treated with or without 500 μM IAA for 5 days prior to harvest. Total RNA was isolated using Illustra RNAspin Mini (GE Healthcare), according to the manufacturer’s protocol. Quantitative PCR was performed using a One Step TB Green PrimeScript RT–PCR Kit II (Takara, #RR086A) on a LightCycler 2.0 instrument (Roche). The thermal cycle was as follows: Stage 1: reverse transcription reaction, 1 cycle (acquisition mode, none): 42°C for 5 min, 95°C for 10 s (temperature shift, 20°C/s). Stage 2: real-time PCR, 40 cycles (acquisition mode, single): 95°C for 5 s, 60°C for 20 s, 72°C for 30 s (temperature shift, 20°C/s). Stage 3: melting curve analysis, 1 cycle (acquisition mode, continuous): 95°C for 0 s, 65°C for 15 s (temperature shift, 20°C/s), followed by 95°C for 0 s (temperature shift, 0.1°C/s). The primers used in this experiment are listed in Table S2. All data were normalized to GAPDH (as a reference).

## Acknowledgements

We thank Dr Katsunori Tanaka for critical reading and Drs Tatsuya Nishino, Tatsuro Takahashi and Hiroyuki Sasanuma for discussion. We also thank Ms. Akemi Mizuguchi, Tomoko Ashikawa and Tomoko Suzuki for technical assistance. This work was supported by JSPS KAKENHI Grants (18H02170 and 18H04719) and by research grants from JST A-STEP (grant number, AS2915150U), the Canon Foundation, the Asahi Glass Foundation and the Takeda Science Foundation.

## Author contributions

VS and MTK designed all experiments. All data were produced by VS. The manuscript was written by VS and MTK.

## References

Bochman ML, Schwacha A. 2009. The Mcm complex: unwinding the mechanism of a replicative helicase. Microbiol Mol Biol Rev 73: 652–683.

Chong JP, Mahbubani HM, Khoo CY, Blow JJ. 1995. Purification of an MCM-containing complex as a component of the DNA replication licensing system. Nature 375: 418–421.

Ding L, Forsburg SL. 2011. Schizosaccharomyces pombe minichromosome maintenance-binding protein (MCM-BP) antagonizes MCM helicase. J Biol Chem 286: 32918–32930.

Edwards MC, Tutter AV, Cvetic C, Gilbert CH, Prokhorova TA, Walter JC. 2002. MCM2-7 complexes bind chromatin in a distributed pattern surrounding the origin recognition complex in Xenopus egg extracts. J Biol Chem 277: 33049–33057.

Ge XQ, Jackson DA, Blow JJ. 2007. Dormant origins licensed by excess Mcm2-7 are required for human cells to survive replicative stress. Genes Dev 21: 3331–3341.

Hahle A, Merz S, Meyners C, Hausch F. 2019. The Many Faces of FKBP51. Biomolecules 9.

Hart T, Chandrashekhar M, Aregger M, Steinhart Z, Brown KR, MacLeod G, Mis M, Zimmermann M, Fradet-Turcotte A, Sun S et al. 2015. High-Resolution CRISPR Screens Reveal Fitness Genes and Genotype-Specific Cancer Liabilities. Cell 163: 1515–1526.

Hills SA, Diffley JF. 2014. DNA replication and oncogene-induced replicative stress. Curr Biol 24: R435–444.

Ibarra A, Schwob E, Mendez J. 2008. Excess MCM proteins protect human cells from replicative stress by licensing backup origins of replication. Proc Natl Acad Sci U S A 105: 8956–8961.

Jagannathan M, Sakwe AM, Nguyen T, Frappier L. 2012. The MCM-associated protein MCM-BP is important for human nuclear morphology. J Cell Sci 125: 133–143.

Kawabata T, Luebben SW, Yamaguchi S, Ilves I, Matise I, Buske T, Botchan MR, Shima N. 2011. Stalled fork rescue via dormant replication origins in unchallenged S phase promotes proper chromosome segregation and tumor suppression. Mol Cell 41: 543–553.

Kim HS. 2019. Genome-wide function of MCM-BP in Trypanosoma brucei DNA replication and transcription. Nucleic Acids Res 47: 634–647.

Kim HS, Park SH, Gunzl A, Cross GA. 2013. MCM-BP is required for repression of life-cycle specific genes transcribed by RNA polymerase I in the mammalian infectious form of Trypanosoma brucei. PLoS One 8: e57001.

Kimura H, Ohtomo T, Yamaguchi M, Ishii A, Sugimoto K. 1996. Mouse MCM proteins: complex formation and transportation to the nucleus. Genes Cells 1: 977–993.

Kubota Y, Mimura S, Nishimoto S, Takisawa H, Nojima H. 1995. Identification of the yeast MCM3-related protein as a component of Xenopus DNA replication licensing factor. Cell 81: 601–609.

Kunnev D, Rusiniak ME, Kudla A, Freeland A, Cady GK, Pruitt SC. 2010. DNA damage response and tumorigenesis in Mcm2-deficient mice. Oncogene 29: 3630-3638.

Kusunoki S, Ishimi Y. 2014. Interaction of human minichromosome maintenance protein-binding protein with minichromosome maintenance 2-7. FEBS J 281:1057-1067.

Leone G, DeGregori J, Yan Z, Jakoi L, Ishida S, Williams RS, Nevins JR. 1998. E2F3 activity is regulated during the cell cycle and is required for the induction of S phase. Genes Dev 12: 2120–2130.

Li JJ, Schnick J, Hayles J, MacNeill SA. 2011. Purification and functional inactivation of the fission yeast MCM(MCM-BP) complex. FEBS Lett 585: 3850–3855.

Lukas C, Savic V, Bekker-Jensen S, Doil C, Neumann B, Pedersen RS, Grofte M, Chan KL, Hickson ID, Bartek J et al. 2011. 53BP1 nuclear bodies form around DNA lesions generated by mitotic transmission of chromosomes under replication stress. Nat Cell Biol 13: 243–253.

Lutzmann M, Grey C, Traver S, Ganier O, Maya-Mendoza A, Ranisavljevic N, Bernex F, Nishiyama A, Montel N, Gavois E et al. 2012. MCM8- and MCM9-deficient mice reveal gametogenesis defects and genome instability due to impaired homologous recombination. Mol Cell 47: 523–534.

Maric M, Maculins T, De Piccoli G, Labib K. 2014. Cdc48 and a ubiquitin ligase drive disassembly of the CMG helicase at the end of DNA replication. Science 346: 1253596.

Masai H, Matsumoto S, You Z, Yoshizawa-Sugata N, Oda M. 2010. Eukaryotic chromosome DNA replication: where, when, and how? Annu Rev Biochem 79: 89–130.

Moreno SP, Bailey R, Campion N, Herron S, Gambus A. 2014. Polyubiquitylation drives replisome disassembly at the termination of DNA replication. Science 346: 477–481.

Natsume T, Kiyomitsu T, Saga Y, Kanemaki MT. 2016. Rapid Protein Depletion in Human Cells by Auxin-Inducible Degron Tagging with Short Homology Donors. Cell Rep 15: 210–218.

Nguyen T, Jagannathan M, Shire K, Frappier L. 2012. Interactions of the human MCM-BP protein with MCM complex components and Dbf4. PLoS One 7: e35931.

Nishimura K, Fukagawa T, Takisawa H, Kakimoto T, Kanemaki M. 2009. An auxin-based degron system for the rapid depletion of proteins in nonplant cells. Nat Methods 6: 917–922.

Nishimura K, Ishiai M, Horikawa K, Fukagawa T, Takata M, Takisawa H, Kanemaki MT. 2012. Mcm8 and Mcm9 form a complex that functions in homologous recombination repair induced by DNA interstrand crosslinks. Mol Cell 47: 511–522.

Nishiyama A, Frappier L, Mechali M. 2011. MCM-BP regulates unloading of the MCM2-7 helicase in late S phase. Genes Dev 25: 165–175.

Ohtani K, Iwanaga R, Nakamura M, Ikeda M, Yabuta N, Tsuruga H, Nojima H. 1999. Cell growth-regulated expression of mammalian MCM5 and MCM6 genes mediated by the transcription factor E2F. Oncogene 18: 2299–2309.

Park J, Long DT, Lee KY, Abbas T, Shibata E, Negishi M, Luo Y, Schimenti JC, Gambus A, Walter JC et al. 2013. The MCM8-MCM9 complex promotes RAD51 recruitment at DNA damage sites to facilitate homologous recombination. Mol Cell Biol 33: 1632–1644.

Pruitt SC, Bailey KJ, Freeland A. 2007. Reduced Mcm2 expression results in severe stem/progenitor cell deficiency and cancer. Stem Cells 25: 3121–3132.

Quimbaya M, Raspe E, Denecker G, De Craene B, Roelandt R, Declercq W, Sagaert X, De Veylder L, Berx G. 2014. Deregulation of the replisome factor MCMBP prompts oncogenesis in colorectal carcinomas through chromosomal instability. Neoplasia 16: 694–709.

Ran FA, Hsu PD, Wright J, Agarwala V, Scott DA, Zhang F. 2013. Genome engineering using the CRISPR-Cas9 system. Nat Protoc 8: 2281–2308.

Sakwe AM, Nguyen T, Athanasopoulos V, Shire K, Frappier L. 2007. Identification and characterization of a novel component of the human minichromosome maintenance complex. Mol Cell Biol 27: 3044–3055.

Santosa V, Martha S, Hirose N, Tanaka K. 2013. The fission yeast minichromosome maintenance (MCM)-binding protein (MCM-BP), Mcb1, regulates MCM function during prereplicative complex formation in DNA replication. J Biol Chem 288: 6864–6880.

Shima N, Alcaraz A, Liachko I, Buske TR, Andrews CA, Munroe RJ, Hartford SA, Tye BK, Schimenti JC. 2007. A viable allele of Mcm4 causes chromosome instability and mammary adenocarcinomas in mice. Nat Genet 39: 93–98.

Taipale M, Tucker G, Peng J, Krykbaeva I, Lin ZY, Larsen B, Choi H, Berger B, Gingras AC, Lindquist S. 2014. A quantitative chaperone interaction network reveals the architecture of cellular protein homeostasis pathways. Cell 158: 434–448.

Takahashi N, Quimbaya M, Schubert V, Lammens T, Vandepoele K, Schubert I, Matsui M, Inze D, Berx G, De Veylder L. 2010. The MCM-binding protein ETG1 aids sister chromatid cohesion required for postreplicative homologous recombination repair. PLoS Genet 6: e1000817.

Todorov IT, Attaran A, Kearsey SE. 1995. BM28, a human member of the MCM2-3-5 family, is displaced from chromatin during DNA replication. J Cell Biol 129: 1433–1445.

Woodward AM, Gohler T, Luciani MG, Oehlmann M, Ge X, Gartner A, Jackson DA, Blow JJ. 2006. Excess Mcm2-7 license dormant origins of replication that can be used under conditions of replicative stress. J Cell Biol 173: 673–683.

Yesbolatova A, Natsume T, Hayashi KI, Kanemaki MT. 2019. Generation of conditional auxin-inducible degron (AID) cells and tight control of degron-fused proteins using the degradation inhibitor auxinole. Methods 164-165: 73–80.

Zeman MK, Cimprich KA. 2014. Causes and consequences of replication stress. Nat Cell Biol 16: 2–9.

