## Supplementary figures and tables for "MCMBP maintains genome integrity by protecting the MCM subunits from degradation"

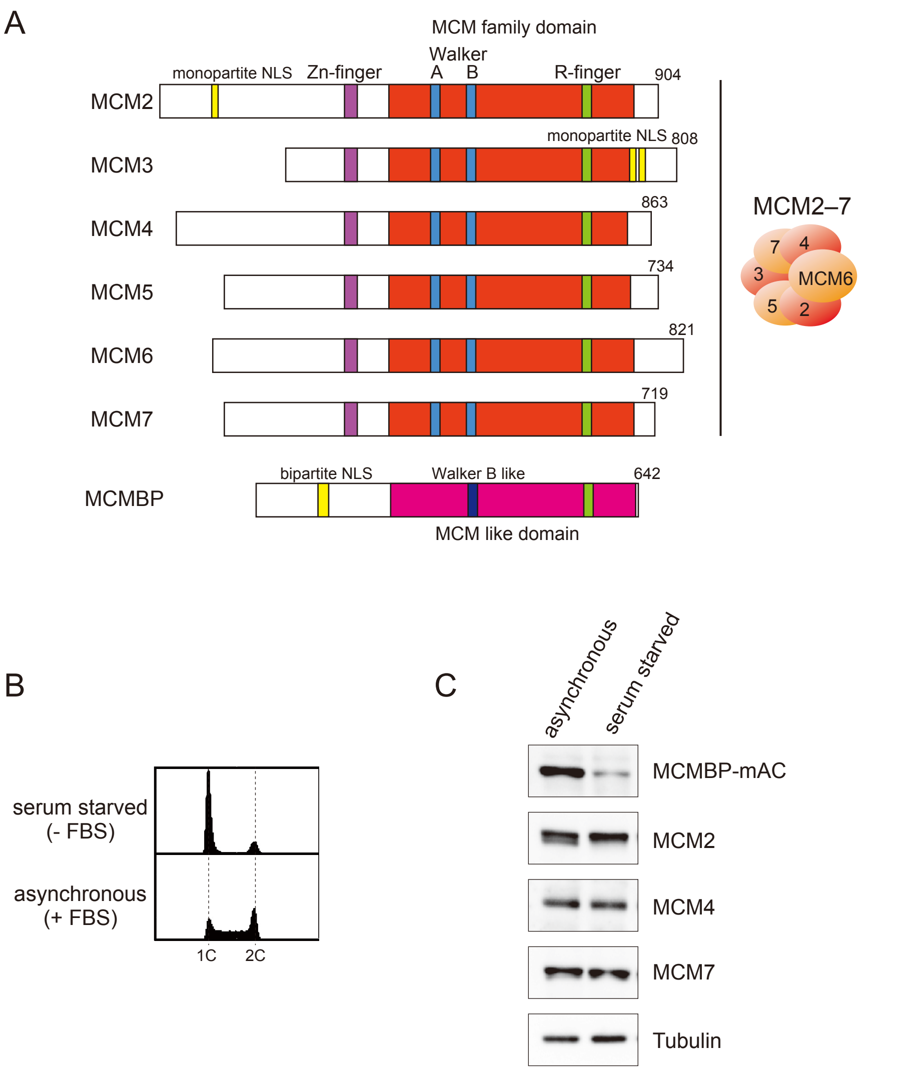
A
MCM family domain
Walker
A
B
monopartite NLS
Zn-finger
R-finger
904
MCM2
monopartite NLS
808
MCM3
863
MCM4
734
MCM5
821
MCM6
719
MCM7
MCM2–7
7
4
3
MCM6
5
2
bipartite NLS
Walker B like
642
MCMBP
MCM like domain
B
serum starved
(- FBS)
asynchronous
(+ FBS)
1C
2C
C
asynchronous
serum starved
MCMBP-mAC
MCM2
MCM4
MCM7
Tubulin

Figure S1

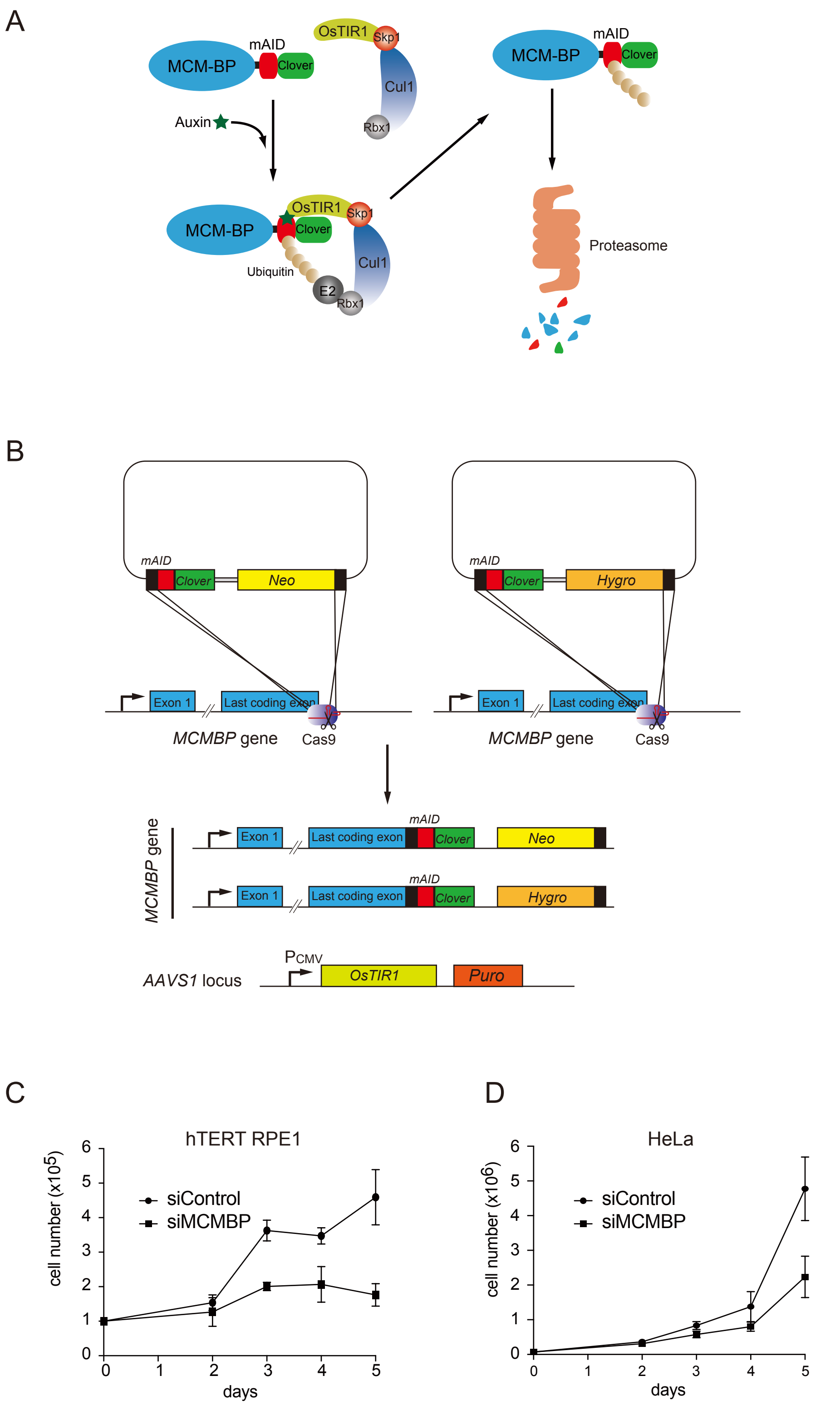
A
MCM-BP
mAID
Clover
OsTIR1
Skp1
Cul1
Rbx1
Auxin
Ubiquitin
E2
Proteasome
B
mAID
Clover
Neo
Hygro
Exon 1
Last coding exon
MCMBP gene
Cas9
AAVS1 locus
PCMV
OsTIR1
Puro
C
hTERT RPE1
cell number (x10^5)
siControl
siMCMBP
days
D
HeLa
cell number (x10^6)
siControl
siMCMBP
days

Figure S2

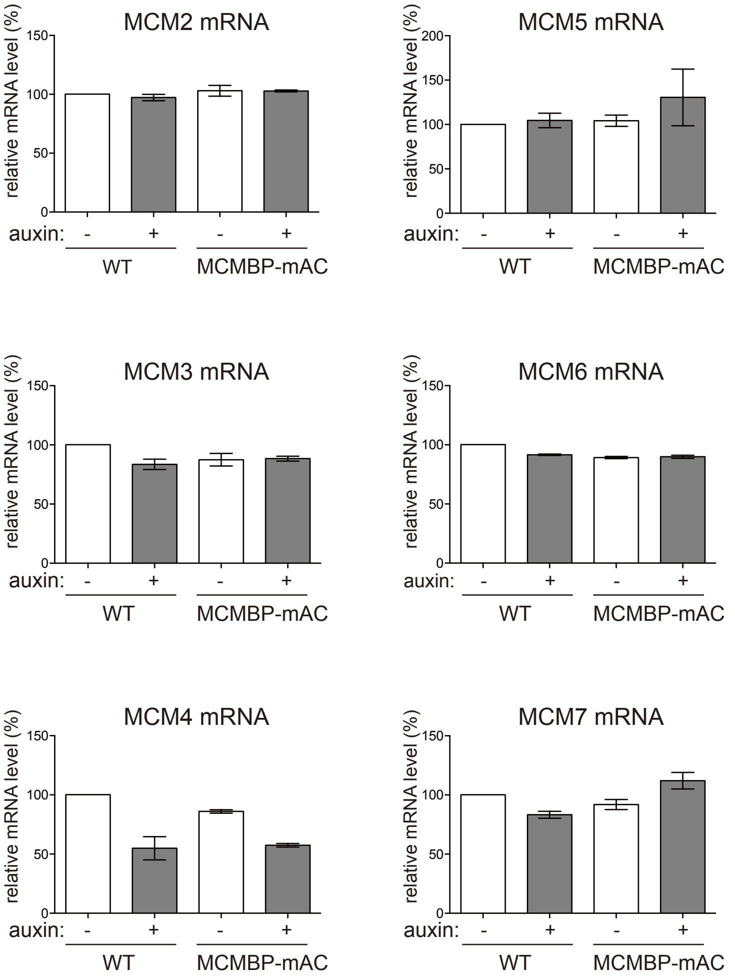
MCM2 mRNA
MCM5 mRNA
MCM3 mRNA
MCM6 mRNA
MCM4 mRNA
MCM7 mRNA
relative mRNA level (%)
0
50
100
150
200
auxin:
-
+
WT
MCMBP-mAC

Figure S3

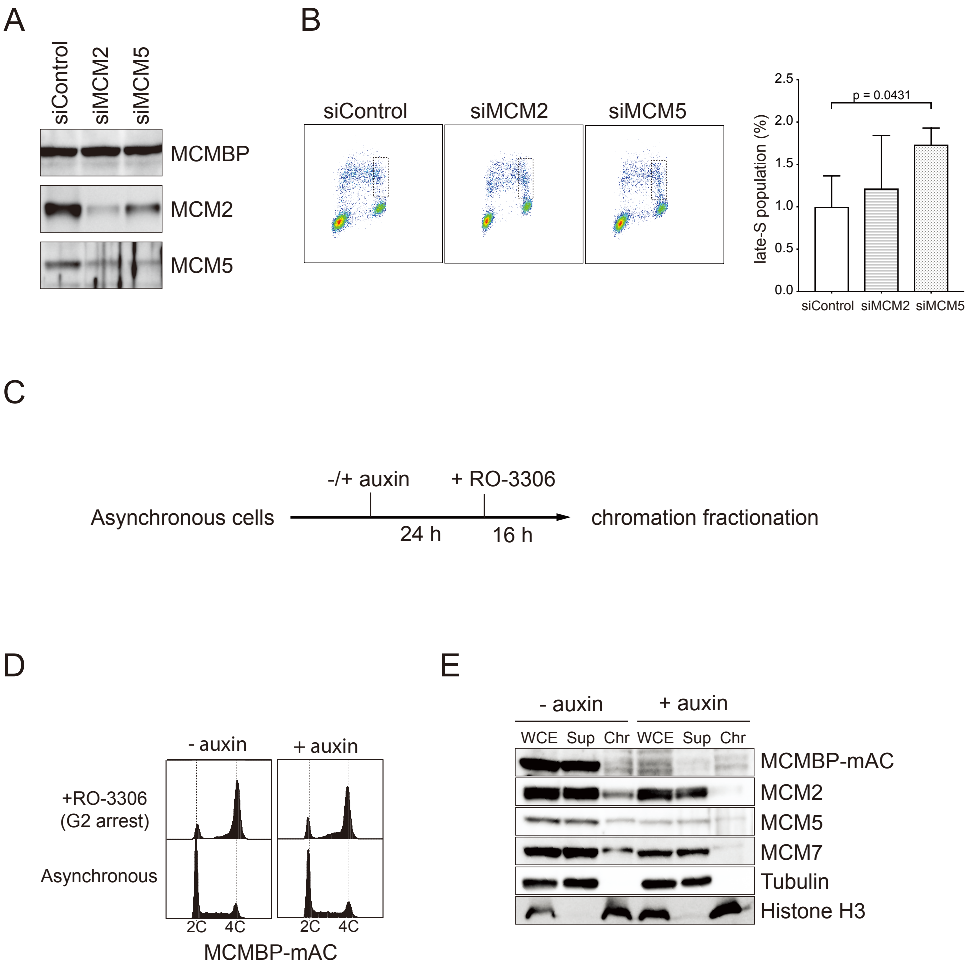
A
siControl
siMCM2
siMCM5
MCMBP
MCM2
MCM5
B
siControl
siMCM2
siMCM5
p = 0.0431
late-S population (%)
2.5
2.0
1.5
1.0
0.5
0.0
siControl
siMCM2
siMCM5
C
-/+ auxin
+ RO-3306
Asynchronous cells
24 h
16 h
chromation fractionation
D
- auxin
+ auxin
+RO-3306
(G2 arrest)
Asynchronous
2C
4C
2C
4C
MCMBP-mAC
E
- auxin
+ auxin
WCE
Sup
Chr
WCE
Sup
Chr
MCMBP-mAC
MCM2
MCM5
MCM7
Tubulin
Histone H3

Figure S4

**Supplementary Information**

**Supplementary Figure Legends**

**Figure S1**: (A) Schematic structure the MCM subunits and MCMBP. Motifs are color coded. Note that MCM2, 3 and MCMBP contains NLS, but the other MCM subunits do not have clear NLS. (B) FACS profile of HCT116 MCMBP-mAC cells arrested by serum starvation. The cells were cultured in medium without FBS for 24 h. (C) Extracts prepared from synchronous and serum starved cells were blotted using the indicated antibodies.

**Figure S2**: (A) Schematic illustration of the Auxin-Inducible Degron (AID) technology to control MCMBP. Endogenous MCMBP was fused with mini-AID–Clover (mAC) at the C-terminus. Auxin binds OsTIR1, which forms the SCF-OsTIR1 E3 ubiquitin ligase, and promotes the interaction between OsTIR1 and mAID. The E3 ubiquitin ligase recruits an E2 ligase resulting polyubiquitylation of the MCMBP-mAC. The fusion protein is rapidly degraded by the proteasome. (B) Construction of the HCT116 MCMBP-mAC cell line. The parental cell expressing OsTIR1 from CMV-OsTIR1 inserted at the AAVS1 locus was used [1]. The mAC tag was inserted using CRISPR–Cas9-based tagging using two donors containing a neomycin or hygromycin resistant marker. After selection by using G418 and hygromycin clones were isolated to confirm the tagging by genomic PCR and Western blotting. (C) The number of siMCMBP-treated hTERT RPE1 cells were counted and plotted. Error bars indicate standard error (n = 3). (D) The number of siMCMBP-treated HeLa cells were counted and plotted. Error bars indicate standard error (n = 3).

**Figure S3**: MCMBP depletion does not affect the mRNA level of the MCM genes. HCT116 MCMBP-mAC cells were treated with or without 500 $\mu$M IAA for 5 days prior to RNA isolation. Error bars indicate standard error (n = 3).

**Figure S4**: (A) HCT116 MCMBP-mAC cells was treated with siMCM2 or siMCM5 for 3 days. Expression of the indicated proteins were tested by Western blotting. (B) siMCM2 or siMCM5 treated cells were analysed by flowcytometry as in Figure 4D. Quantified population of cells in late S phase was shown in right. Error bars indicate standard error (n = 3). (C) Experimental scheme to arrest HCT116 MCMBP-mAC cells in G2 phase. (D) G2 arrest was confirmed by flowcytometry. (E) MCMBP depletion does not affect dissociation of the MCM2–7 from chromatin DNA. G2 arrested cells were harvested and fractionated to test chromatin association of the indicated proteins.

**Table S1**

| **Primary antibody** | **Manufacture** | **Catalogue number** |
|---|---|---|
| Rabbit polyclonal MCMBP | PGI Proteintech | 19573-1-AP |
| Mouse monoclonal mini AID (mAID) | MBL | M214-3 |
| Goat polyclonal hMCM2 (N-19) | SantaCruz | sc-9839 |
| Mouse monoclonal hMCM4 (G-7) | SantaCruz | sc-28317 |
| Rabbit polyclonal hMCM5 (H-300) | SantaCruz | sc-22780 |
| Mouse monoclonal hMCM7 | MBL | M049-3 |
| Mouse monoclonal α-tubulin | MBL | M175-3 |
| Goat polyclonal Histone H3 (C-16) | SantaCruz | sc-8654 |
| Mouse monoclonal RFP | MBL | M204-3 |
| Mouse monoclonal anti-Flag M2 | Sigma | F1804-5MG |
| Rabbit polyclonal 53BP1 (H-300) | SantaCruz | sc-22760 |
| Mouse monoclonal BrdU | BD | 347580 |

| **Secondary antibody** | **Manufacture** | **Catalogue number** |
|---|---|---|

| | | |
|---|---|---|
| HRP-conjugated goat polyclonal Rabbit-IgG | GE Healthcare | NA934 |
| HRP-conjugated goat polyclonal Mouse-IgG | SantaCruz | PI-2000 |
| HRP-conjugated donkey polyclonal Goat-IgG | SantaCruz | sc-2020 |
| Goat polyclonal StarBright Blue 700 Anti-Rabbit IgG | BioRad | 12004161 |
| Goat polyclonal StarBright Blue 700 Anti-Mouse IgG | BioRad | 12004158 |
| Goat polyclonal StarBright Blue 520 Anti-Mouse IgG | BioRad | 12005866 |
| Goat polyclonal StarBright Blue 520 Anti-Rabbit IgG | BioRad | 12005869 |
| hFAB Rhodamine anti-tubulin IgG | BioRad | 12004165 |
| Alexa Fluor 488-conjugated goat polyclonal anti-Rabbit IgG | Life Technologies | A11034 |
| Alexa Fluor 594-conjugated goat polyclonal anti-Mouse IgG | Life Technologies | A11032 |
| Goat polyclonal FITC-conjugated anti-mouse IgG | Jackson Laboratory | 115-095-164 |

### Table S2

| **Oligonucleotides** | |
|---|---|
| qPCR primers for GAPDH | FW: GGTGTGAACCATGAGAAGTATGA |
| | RV: GAGTCCTTCCACGATACCAAAG |
| qPCR primers for MCM2 | FW: CTGAGAAGGACTTGGTGGATAAG |
| | RV: CCTGAAGAGCTCACTGTCATAAA |
| qPCR primers for MCM3 | FW: CTATGCCAAGCAGTATGAGGAG |
| | RV: TGAGGAAGCAGGAGGTAAGA |
| qPCR primers for MCM4 | FW: CCTCATTGGTAAAGGGCTAGAG |
| | RV: TAGCCAGGGTGACAGAGTAA |
| qPCR primers for MCM5 | FW: CCTTCGTCCCGGAATTTCAT |
| | RV: CGGTCATCTTCTCGCATCTT |
| qPCR primers for MCM6 | FW: TACCAGCCTGTTCCCTACTAT |
| | RV: GTCCCTTCTCCTGTTGTCTTT |
| qPCR primers for MCM7 | FW: TAGACAAGAAGACGGCGAAAG |
| | RV: GGAGGAGAATCGCTCTTAAAGG |

### References

1. Natsume, T., Kiyomitsu, T., Saga, Y., and Kanemaki, M.T. (2016). Rapid Protein Depletion in Human Cells by Auxin-Inducible Degron Tagging with Short Homology Donors. Cell Rep *15*, 210-218.
